# TRAIL pathway suppression of cancer cell growth and immune cell-mediated tumor cell-killing in a senescent fibroblast-constructed tumor microenvironment

**DOI:** 10.1101/2023.11.30.569479

**Authors:** Shengliang Zhang, Kelsey E. Huntington, Lanlan Zhou, Attila A. Seyhan, Bianca Kun, Benedito A. Carneiro, Jill Kreiling, John M. Sedivy, Wafik S. El-Deiry

## Abstract

Cellular senescence and the associated secretory phenotype (SASP) promote cancer in the aging population. During aging or upon chemotherapy exposure, cellular and molecular changes occur in non-cancerous cells and alter responses to cancer therapy, primarily via modifications in the tumor microenvironment (TME) and immune response. Targeting senescent cells through removal, modulation of the SASP, or cellular reprogramming represent promising therapeutic avenues for treating cancer. We elucidate an interplay between cancer cells, immune cells, and senescent fibroblasts and describe the impact of fibroblast senescence on tumor growth and response to cancer therapy. Cytokine profiling reveals dynamic changes in SASP production during etoposide-induced senescence in IMR90 fibroblasts. We show that SASP is partially regulated by p21 (WAF1; CDKN1A), leading to the downregulation of anti-tumorigenic cytokines and upregulation of pro-tumorigenic cytokines. Senescent fibroblasts promote bystander cancer cell growth via a p21-driven SASP. These results provide strategies to target the p21-driven SASP in the TME during cancer therapy. Treatment with TRAIL or TRAIL-inducing Dordaviprone (TIC10/ONC201) reduces cell viability of tumor cells co-cultured with senescent or proliferating fibroblasts and promotes immune-mediated tumor cell-killing in co-culture with senescent IMR90 fibroblasts. ONC201 combined with senolytic drugs (e.g., Navitoclax, Lamivudine) synergizes towards tumor suppression. These results indicate that senolytic therapies may be combined with cancer therapies to target senescence-associated changes in the TME including for modulation of the senescent cytokine landscape.

## Introduction

Age is a risk factor for cancer and cancer is a disease of aging [1]. This is a highly relevant risk factor as more than two-thirds of new cancer cases are diagnosed in individuals over the age of 60 [2]. One key component of aging is cellular senescence, a form of terminal cell cycle arrest that includes a <u>S</u>enescence-<u>A</u>ssociated <u>S</u>ecretory <u>P</u>henotype (SASP). Although cellular senescence leads to tumor suppression, there is ample evidence that persistently senescent cells acquire pro-tumorigenic properties and eventually contribute to cancer progression [3]. Moreover, anticancer therapies including chemotherapy, radiation or PARP inhibitors induce cellular senescence, a phenomenon termed therapy-induced senescence [4].

Accumulation of cellular senescence contributes to a tumor permissive microenvironment and a dampened immune response [5–7]. This is partially due to the SASP and the resulting modification in cytokine, chemokine, and growth factor profiles by the senescent cells [8]. Frequently observed SASP proteins include cytokines interleukin 1 β (IL-1β), interleukin 6 (IL-6), interleukin 8 (IL-8), and granulocyte-macrophage colony-stimulating factor (GM-CSF), and chemokines macrophage inflammatory protein 1 alpha (MIP-1 alpha), monocyte chemoattractant protein-1 (MCP-1), regulated upon activation, normal T Cell expressed and secreted (RANTES/CCL5), and growth-regulated oncogene-alpha (GRO alpha) [8, 9]. Senescence-associated modifications in cytokine and chemokine profiles have the potential to modify immune cell recruitment and activation, among other diverse cellular responses [10]. Therapeutic treatment of senescent cells may differentially modify cytokine profiles. The cytokine landscapes of senescent cells need to be characterized for therapeutic targeting of cellular senescence during and after cancer treatment.

Accordingly, emerging therapeutic opportunities include 1) clearance of senescent cells by utilizing the host’s immune system or 2) combination treatment with senolytic agents [11]. Here, we utilize senolytic Bcl-2 inhibitor ABT263 (Navitoclax), anti-retroviral nucleoside analog 3TC (Lamivudine), TRAIL/DR5-inducing ClpP agonists Dordaviprone (TIC10/ONC201), and its potent analogue ONC212. Senolytic agents and TRAIL pathway agonists may promote anti-tumor immune responses in a senescent TME and have implications for response to cancer therapies, especially immune-based therapies or emergent resistance mechanisms after chemotherapy, radiation or targeted therapies [11–13].

## Methods

### Cell culture

Cell lines used in this study include human colorectal adenocarcinoma HT-29 cells (ATCC), normal lung fibroblast IMR90 cells (ATCC), human CD8+ cytotoxic TALL-104 cells (ATCC), and human natural killer NK-92 cells (previously provided by Dr. Kerry Campbell at Fox Chase Cancer Center). HT-29 cells were cultured in McCoy’s 5A (modified) Medium, IMR90 cells were cultured in Eagle’s Minimal Essential Medium, and TALL-104 cells were cultured in RPMI 1640 Medium. All cell line media was supplemented with 10% FBS and 1% Penicillin-Streptomycin, except for TALL-104 cells which were supplemented with 20% FBS. NK-92 cells were cultured in Alpha Minimum Essential medium without ribonucleosides and deoxyribonucleosides but with 2 mM L-glutamine and 1.5 g/L sodium bicarbonate supplemented with 0.2 mM inositol; 0.1 mM 2- mercaptoethanol; 0.02 mM folic acid; 12.5% horse serum and 12.5% FBS. Recombinant human IL-2 (Miltenyi cat. no. 130–097744) with a final concentration of 100 units/mL was added to the media of the NK-92 and TALL-104 cell lines. All cell lines were incubated at 37°C in a humidified atmosphere containing 5% CO2. Cell lines were authenticated and tested to ensure the cultures were free of mycoplasma infection.

### Senescence induction

Senescent IMR90s were induced as previously reported [14]. Briefly, approximately 0.5 x 10^6^ human fibroblast IMR90 cells at early passages were seeded on 10 cm plates and treated with etoposide (40 μM) for 72 hours, after which cells were cultured with etoposide-free completed culture medium (EMEM supplemented with 10% FBS) and the culture media were changed every three days for three weeks.

### Co-culture assay

Tumor cells and fibroblasts were plated and allowed to grow for 24 hours in their culture media before the addition of immune cells (NK-92 or TALL-104). Cells were labeled with either CellTracker Green CMFDA or CellTracker Blue CMAC dyes (Invitrogen, Waltham, MA). Red fluorescent ethidium homodimer (EthD-1) was added at the beginning of the tri-culture to detect dead cells (Invitrogen, Waltham, MA). For the quantification of dead/live cells, fluorescence microscopy was used to take images at 10x magnification. The number of red/green color cells in random fields was determined using thresholding and particle analysis in the Fiji [15] modification of ImageJ and expressed as a dead/live cell ratio.

### Collection of cell supernatant samples for cytokine profiling

Cells were plated at 3.5 × 10^4^ cells in a 48-well plate (Thermo Fisher Scientific, Waltham, MA, USA) in complete medium and incubated at 37°C with 5% CO2. At 24 hours after plating, almost all the tumor cells were adherent to the bottom of the wells and the complete medium was removed and replaced with the drug-containing medium. After 48 hours of treatment, cell culture supernatants were collected and centrifuged to remove cellular debris, Samples were frozen at - 80°C for measurement of cytokines, chemokines, and growth factors in batch.

### Cytokine profiling for drug treatment experiments

An R&D systems Human Premixed Multi-Analyte Kit (R&D Systems, Inc., Minneapolis, MN) was run on a Luminex 200 Instrument (LX200-XPON-RUO, Luminex Corporation, Austin, TX) according to the manufacturer’s instructions. Cell culture supernatant levels of TNF-alpha, IL-6, IL-8/CXCL8, Ferritin, IFN-beta, IL-10, CCL2/JE/MCP-1, VEGF, CXCL13/BLC/BCA-1, IFN-gamma, CCL20/MIP-3 alpha, CCL3/MIP-1 alpha, CCL22/MDC, CCL4/MIP-1 beta, IL-4, IL-17/IL- 17a, TRAIL R2/TNFRSF10B, GM-CSF, CXCL5/ENA-78, CXCL9/MIG, G-CSF, CXCL11/I-TAC,Granzyme B, CCL5/RANTES, Prolactin, IFN-alpha, CXCL14/BRAK, IL-12/IL-23 p40, CXCL10/IP- 10/CRG2, CCL7/MCP-3/MARC, IL-7, CCL8/MCP-2, TRANCE/TNFSF11/RANK L, IL-15, TRAIL R3/TNFRSF10C, CCL11/Eotaxin, IL-18/IL-1F4, TRAIL/TNFSF10, IL-21, and C-Reactive Protein/CRP were measured for the drug treatment experiments. Analyte values were reported in picograms per milliliter (pg/mL).

### Cytokine profiling for senescence induction experiments

An R&D systems Human Premixed Multi-Analyte Kit (R&D Systems, Inc., Minneapolis, MN) was run on a Luminex 200 Instrument (LX200-XPON-RUO, Luminex Corporation, Austin, TX) according to the manufacturer’s instructions. Cell culture supernatant levels of CCL2/JE/MCP-1, CCL5/RANTES, CCL7/MCP-3/MARC, CXCL1/GRO alpha/KC/CINC-1, CXCL2/GRO beta/MIP- 2/CINC-3, CXCL10/IP-10/CRG-2, G-CSF, GM-CSF, IFN-gamma, IGFBP-3, IL-1 alpha/IL-1F1, IL-1 beta/IL-1F2, IL-1ra/IL-1F3, IL-6, IL-8/CXCL8, IL-12/IL-23 p40, M-CSF, MMP-1, MMP-2, MMP-9, Serpin E1/PAI-1, SPARC, TNF-alpha, TRAIL/TNFSF10 were measured for the induction of senescence experiments. Analyte values were reported in picograms per milliliter (pg/mL).

### Single-cell cytokine profiling assay

Cells were prepared according to the protocol and loaded onto an IsoCode chip coated with multiplexed antibodies which allows detection of 32 cytokines at a single-cell level using IsoPlexis Human Adaptive Immune Chips and IsoPlexis technology.

### Western blot analysis

Cells were lysed in RIPA buffer (Sigma-Aldrich, St. Louis, MO, USA) containing cocktail protease inhibitors (Roche, Basel, Switzerland). Equal amounts of cell lysates were electrophoresed through 4-12% SDS-PAGE then transferred to PVDF membranes. The transferred PVDF membranes were blocked with 5% skim milk at room temperature, then incubated with primary antibodies incubated in a blocking buffer at 4°C overnight. Antibody binding was detected on PVDF with appropriate HRP-conjugated secondary antibodies by a Syngene imaging system (Syngene, Bangalore, India).

### Colony formation assay

A total of 300 cancer cells per well were co-cultured with the same number of senescent IMR90 or proliferating IMR90 cells in 12-well plates. Cells were treated with different drugs for three days, then cells were cultured with drug-free complete medium for two weeks with fresh medium being changed every three days. Cell colonies were fixed with 10% formalin and stained with 0.05% crystal violet at the end of the experiments.

### Knockdown of Gene expression by siRNA transient transfection

Cells were seeded in 96-well plate and transfected with siRNA using lipofectamine RNAi MAX (Invitrogen) as described in the manufacturer’s protocol. At 24 hours after siRNA transfection, cells were co-cultured with cancer cells carrying a luciferase reporter for three days.

### Luciferase reporter assay

Cancer cells carrying a constitutive luciferase reporter were co-cultured with senescent or proliferating IMR90 cells. The luciferase reporter expression in cancer cells was examined based on bioluminescence using the IVIS imaging system (PerkinElmer, Hopkin, MA, USA).

### SAβ-gal staining

IMR90 cells seeded on the plate were fixed and stained for SA-β-gal activity with the senescence β-Galactosidase Staining Kit according to the protocol (Cell Signaling, #9860S).

### In vivo experiments

One million HT29 carrying a constitutive luciferase reporter (HT29-luc) cancer cells with the same amount of senescent or proliferating IMR90 cells in a 200 μl suspension of 1:1 Matrigel (BD) were implanted subcutaneously in the flanks of nude mice (female, 4-6 weeks old, Taconic Biosciences). Mice were treated with ABT263 (in DMSO:polyethylene glycol 400:Phosal 50 PG, gavage at 50 mg per kg body weight per day (mg/kg/d) for 10 days. TIC10/ONC201 (gavage at 50 mg per kg body weight weekly), or the combination when the xenografted tumor reached a volume of ∼200 mm^3^ (day 0 for the treatment). Tumors were measured twice a week. Bioluminescence imaging was performed on each mouse using IVIS imager weekly.

### Statistical analysis

The statistical significance of differences between pairs was determined using unpaired Student’s t tests. The statistical significance between groups was determined using a One-way Anova followed by a post-hoc Tukey’s multiple comparisons test. The minimal level of significance was P < 0.05. The following symbols * and ** represent, P < 0.05 and P < 0.01, respectively.

## Results

### Modulation of fibroblast cytokine profiles during chemotherapy-induced senescence

We first sought to characterize the changes in cytokine profiles that IMR90 fibroblasts undergo during the induction of senescence. IMR90 fibroblasts were treated with etoposide to induce senescence, and cell culture supernatant samples were collected before treatment, one-week post-etoposide, and three-weeks post-etoposide **(Fig. 1A)**. Etoposide-induced senescent fibroblasts were observed using senescence-associated β-galactosidase (SA-β-gal) staining **(Fig. 1B).** Cell culture supernatants were analyzed using a custom cytokine, chemokine, and growth factor profiling panel that was designed to analyze SASP-mediated changes in pro- and anti-inflammatory mediators of anti-tumor immunity. Several analytes trended down during the induction of senescence including Tumor Necrosis Factor alpha (TNF alpha), Matrix Metallopeptidase 9 (MMP-9), Interferon gamma (IFN-gamma), Interleukin 12 (IL-12), Insulin-like Growth Factor-Binding Protein 3 (IGFBP-3), CCL7, Monocyte Chemotactic Protein-3 (MCP- 3/MARC), and TNF-Related Apoptosis-Inducing Ligand (TRAIL/TNFSF10) **(Fig. 1C).** By contrast, analytes with a positive trend towards upregulation during the induction of senescence included Interleukin 6 (IL-6), Interferon gamma-Induced Protein 10 (CXCL10/IP-10), Monocyte Chemoattractant Protein-1 (CCL2/MCP-1), Interleukin 1 alpha (IL-1 alpha), Granulocyte colony-stimulating factor (G-CSF), Interleukin 8 (IL-8/CXCL8), Macrophage Inflammatory Protein 2-alpha (CXCL2/GRO beta/MIP-2/CINC-3), Matrix Metallopeptidase 1 (MMP-1), Matrix Metallopeptidase (MMP-2), Growth-Regulated Oncogene alpha (CXCL1/GRO alpha/KC/CINC1), and Interleukin 1 beta (IL-1 beta) **(Fig. 1D)**. Cytokinome modifications are consistent with previous reports of SASP secretion by senescent cells, validating the induction of IMR90 fibroblast senescence induced by etoposide.

**Figure 1.**
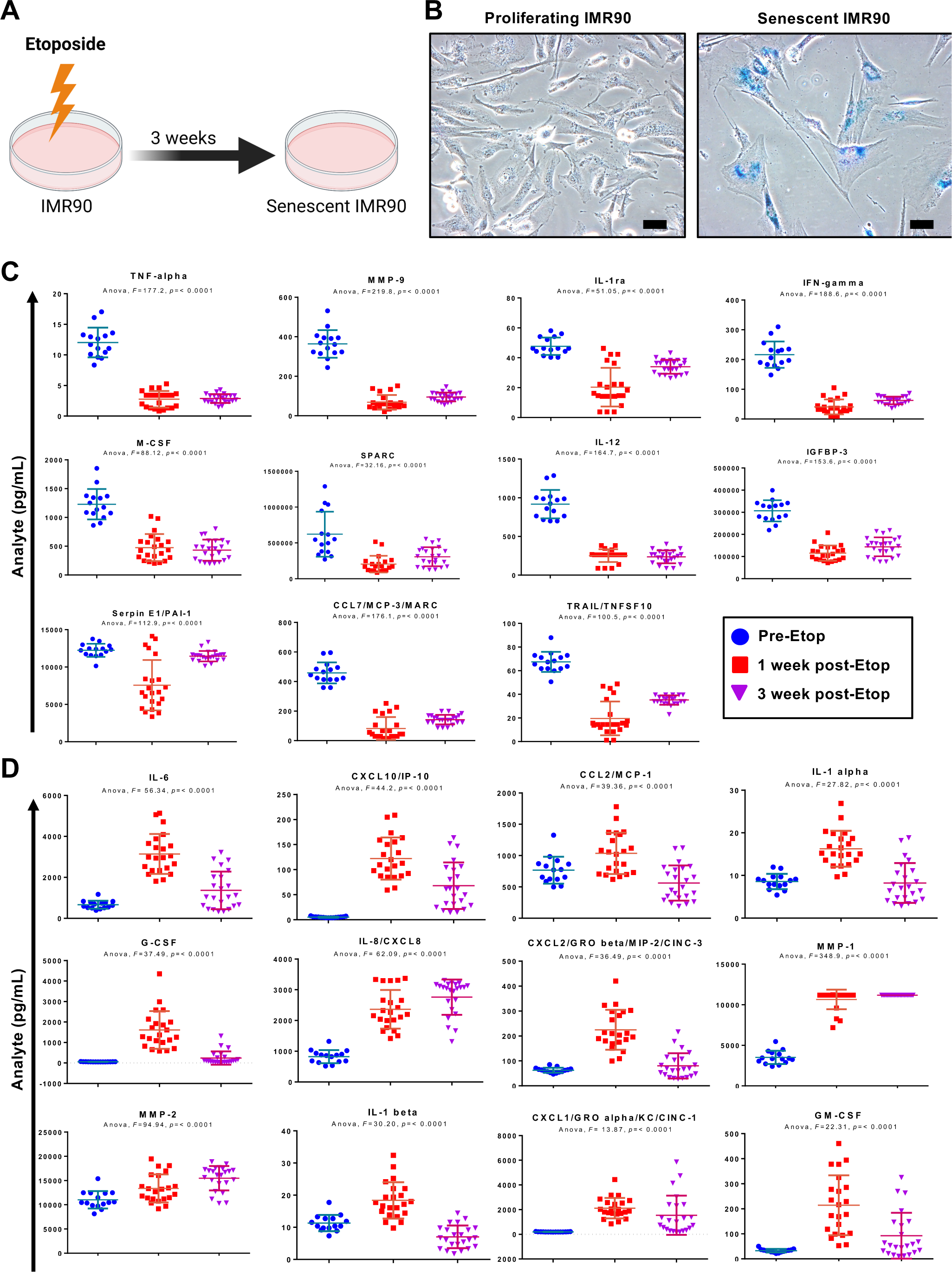
Cytokine profiles are modulated during senescence induction in IMR90 fibroblasts. A. Schema of IMR90 senescence induced by etoposide. B. SA-β-gal staining of senescent IMR90 fibroblasts at 3 weeks post-treatment with etoposide (right panel). 200x magnification, scale bar represents 100 µm. C. Cytokine profiles of analytes that decreased post-etoposide treatment in IMR90 cells. D. Cytokine profiles of analytes that increased post-etoposide treatment in IMR90 cells. Blue circles represent pre-etoposide analyte concentrations in picograms per milliliter (pg/mL), red squares represent analytes concentrations 1-week post-etoposide, and purple inverted triangles represent analyte concentrations 3-weeks post- etoposide.

### Senescent fibroblasts promote tumor cell proliferation in culture and tumor growth *in vivo*

To investigate the effect of senescent fibroblast cells in the TME on tumor growth, we set up an *in vitro* co-culture system in which HT29 colorectal cancer (CRC) cells carrying a constitutive- luciferase reporter (HT29-Luc) were co-cultured with IMR90 fibroblasts **(Fig. 2A**). IMR90 fibroblasts are the most widely used fibroblast senescence model in the research community including for studies of cancer-senescent cell interactions. The production of luciferase by HT29- Luc cancer cells was confirmed using bioluminescence. The same number of HT29-Luc cells were seeded with either senescent or proliferating IMR90 cells and showed similar bioluminescence on day 0 **(Fig. 2B)**. On days 3 and 6 , we detected an increase in bioluminescence in the HT29-Luc cells co-cultured with the senescent IMR90 cells as compared to those co-cultured with the proliferating cells **(Fig. 2B, supplementary figure 1).** The cell growth was further confirmed under microscopy at day 6 by the observation that there were more HT29 cells growing in the presence of the senescent IMR90 cells (**Figure 2C**). These results suggest that cancer cell growth in the presence of senescent IMR90 cells is greater than in the presence of proliferating fibroblasts.

**Figure 2.**
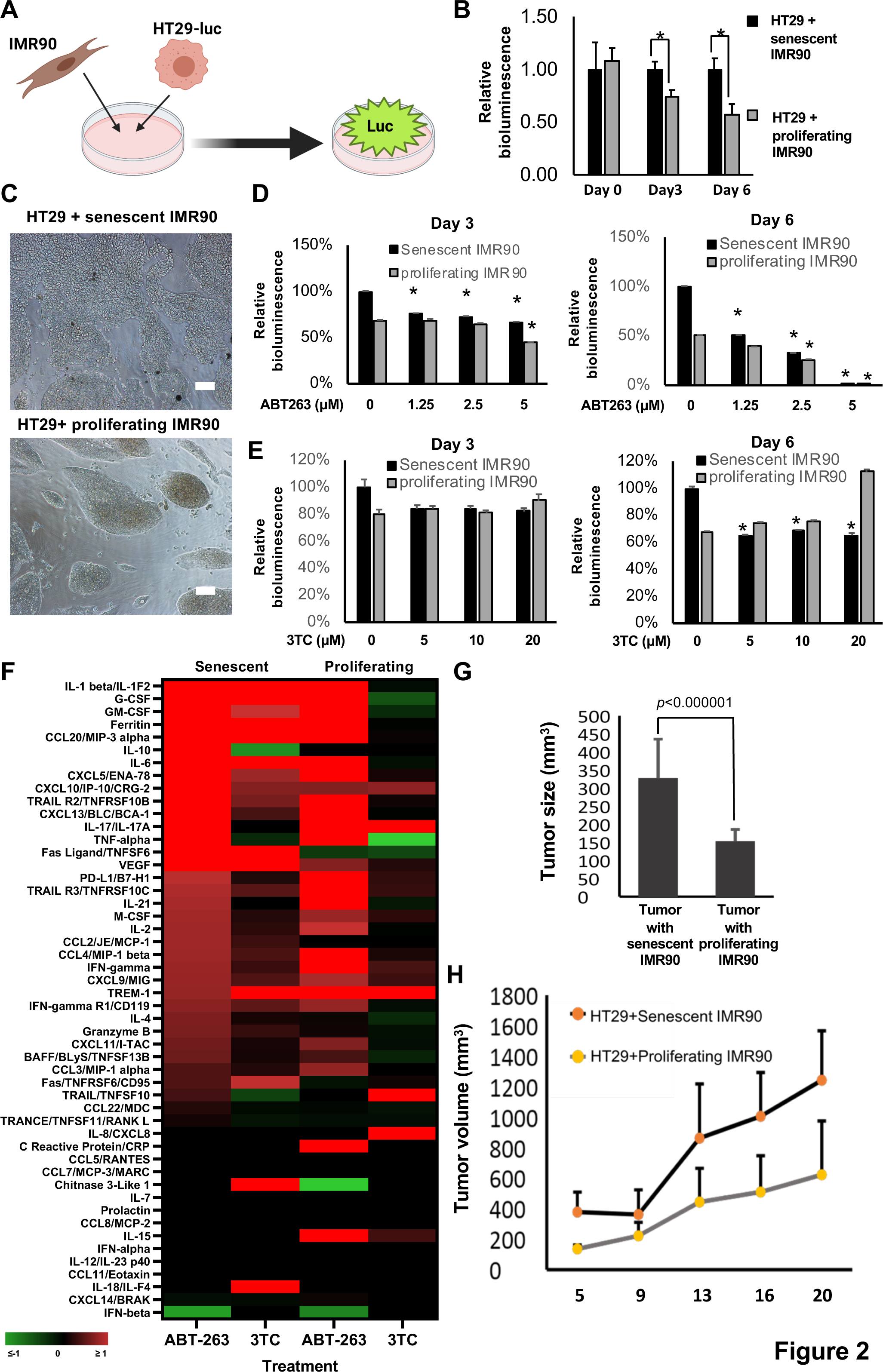
**Senescent fibroblasts promote tumor cell proliferation in co-culture and tumor growth in vivo**.. A. Schema of co-culture system with constitutive Luciferase reporter expression in HT29 cancer cells. B. Relative bioluminescence in a two-cell co-culture system using HT29 tumor cells and IMR90 fibroblasts. Relative bioluminescence was normalized to that of HT29-luc cocultured with senescent IMR90 at each day indicated. C. Imaging of tumor cells in co-culture system at day 6. Scale bar represents 100 microns. D. Bioluminescence in co-culture system at 3 and 6 days post-ABT263 treatment. E. Bioluminescence in co-culture system at 3 and 6 days post-3TC treatment. F. Heat map representing cytokine, chemokine, and growth factor profiles. Fold changes post-treatment with ABT-263 (5µM) or 3TC (10µM) are shown in both senescent and proliferating IMR90s. G. HT29 mouse xenograft average tumor growth volumes on day 5 after HT29 cells were subcutaneously implanted with senescent or proliferating IMR90 cells. H. Tumor growth curves of HT29 CRC xenografts in mice. Tumor growth is reported according to tumor size measurements. Tumor volumes (G and H) were measured by a caliper every three days. Relative bioluminescence was normalized to control in each cohort, respectively. Data are expressed as mean ± SD. *, *p*<0.05.

We attempted to block the effects of celllular senescence by clearance of senescent cells using senolytic drug ABT263 or inhibiting the SASP using the senolytic drug 3TC. Both ABT263 and 3TC significantly reduced bioluminescence of HT29-Luc cells co-cultured with senescent IMR90 cells.**(Fig.2D, 2E).** Treatment with ABT263 reduced the cell viability of senescent fibroblasts without affecting proliferating IMR90 cell viability **(Supplementary Fig. 2A).** This result agrees with previous reports showing the ability of ABT263 to clear senescent cells [16]. Cytokine profiling reveals TNF-alpha, Fas ligand, IFN-γ and TRAIL, but the IFN-γ and TRAIL induction is modest in the senescent IMR90 cells with ABT263 treatment **(Fig. 2F).** By contrast, 3TC did not reduce cell viability of senescent or proliferating IMR90 cells **(Supplementary Fig. 2B).** 3TC is known to downregulate the SASP [17] and we similarly observed downregulation of SASP factors in senescent IMR90 cells after treatment with 3TC **(Fig. 2F),** including IL-10 and TRAIL/TNFSF10 [18, 19]. These results suggest that inhibition of SASP and fibroblast senescence can contribute to tumor suppression in the tumor microenvironment.

We investigated the effect of senescent fibroblasts on tumor formation and growth in the tumor microenvironment (TME) in mice. We generated *in vivo* tumor xenografts by implanting HT29 CRC cells with senescent IMR90 cells in nude mice. Five days after cancer cell implantation with senescent or proliferating IMR90 cells, we observed that tumors in the presence of senescent IMR90 cells were significantly larger than those in the presence of proliferating IMR90 cells **(Fig. 2G)**. These *in vivo* results suggest that senescent IMR90 cells facilitates cancer cell proliferation and tumor formation by modulating the TME. Tumor growth curves showed a faster growth rate and size of tumors in the presence of senescent IMR90s as compared to tumors in the presence of proliferating IMR90s **(Fig. 2H)**.

### p21 (WAF1; CDKN1A) is essential for senescent fibroblast SASP production and bystander cancer cell growth

Chemotherapy induces cancer associated fibroblast (CAF) senescence in which p21(WAF1; CDKN1A), the cyclin-dependent kinase inhibitor, arrests the cell cycle. We explored the effect of host p21 in senescent fibroblasts on bystander cancer cell growth. We transiently knocked-down p21 expression in senescent fibroblasts using siRNA. The knockdown of p21 in the senescent fibroblasts suppressed bystander cancer cell growth. As shown in **Figure 3**, bioluminescence was reduced in bystander cancer cells co-cultured with p21-knockdown senescent fibroblast cells (**Fig. 3A-B**). SA-β-gal staining showed strong SA-β-gal activity around nuclei in senescent fibroblasts with p21-knockdown as compared to cells with siRNA control (**Fig. 3C**). Further cytokine profiling showed two panels of SASP-associated proteins upon p21-knockdown in senescent cells (**Fig. 3D**). We observed decreases of cytokines such as, CHI3L1, CCL2 and IL-8 and IL-2 and increases of cytokines such TNFα, IL-21 and ferritin with p21-knockdown in senescent fibroblasts. These results suggest that p21 drives SASP in senescent fibroblasts, in addition to its function in cell cycle arrest. We further knocked-down TNFα and p21 expression using siRNA in senescent IMR90 cells, and the knockdown of TNFα partially or modestly rescued the anti-proliferation effect of p21-knockdown in senescent IMR90 cells on the bystander cancer cells (**Fig. 3E and F**). These results suggest that senescent fibroblasts promote bystander cancer cell growth via a p21-driven SASP in the TME (**Fig.3G**).

**Figure 3.**
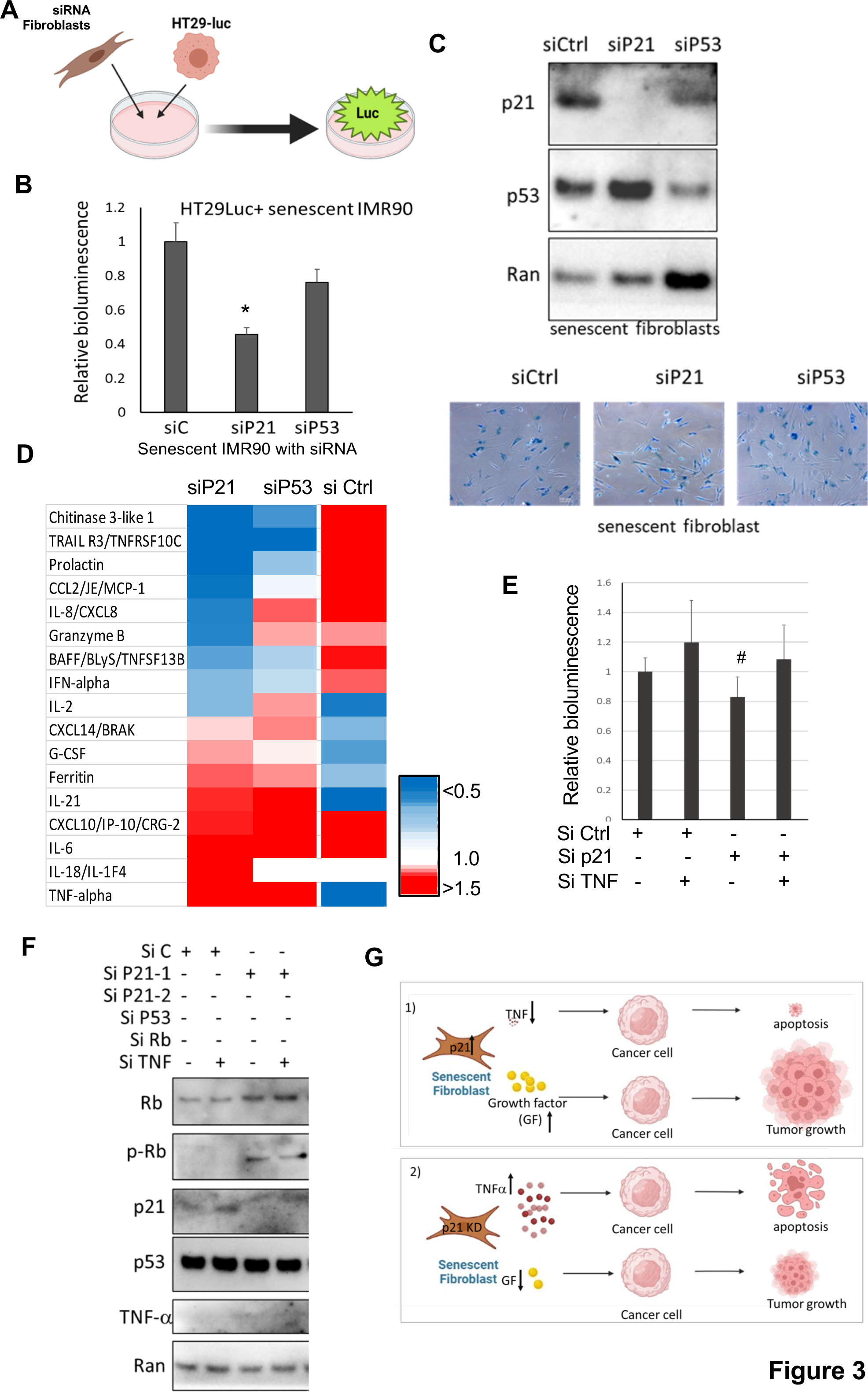
p21-knockdown influences SASP in senescent IMR90 and suppresses bystander cell growth. A. Schema of culture of p21-knockdown senescent IMR90 with bystander cancer cells. B. Bioluminescence of bystander cells in two-cell co-culture system on day 3. C. SA-β-gal staining of p21-Knockdown senescent IMR90 (lower panel). p21 expression in p21-knockdown or p53-knockdown senescent IMR90 cells is shown (upper panel). D. Heat map representing cytokine, chemokine, and growth factor profiles. Fold changes are shown for p21-knockdown senescent IMR90 cells. Relative cytokine levels were normalized to those of si-Ctrl in senescent IMR90 cells. Relative cytokine levels from senescent IMR90 with si-Ctrl were normalized to those from proliferating IMR90 cells. E. Bioluminescence of bystander HT29 cancer cells in two-cell co- culture system with TNFα- knockdown senescent IMR90 cells. Data are expressed as mean ± SD. #, p≤0.1. F. Western blot assay for TNFα and p21 expression in senescent IMR90 cells (E). G. Schematic representation of senescent fibroblasts promote bystander cancer cell growth via p21-driven SASP in the TME. Relative bioluminescence was normalized to control for each cohort, respectively. Data are expressed as mean ± SD. *, *p*<0.05.

### TRAIL and TRAIL-inducing TIC10/ONC201 reduce cancer cell growth mediated by senescent or proliferating fibroblasts

Tumor Necrosis Factor (<u>T</u>NF)-<u>R</u>elated <u>A</u>poptosis-Inducing <u>L</u>igand (TRAIL) is a ligand for death receptor 5 (DR5) and promotes tumor cell death via induction of cellular apoptosis. We noted that secretion of TRAIL and/or shedding of soluble DR5 (sDR5) was reduced during the induction of IMR90 senescence **(Fig. 1C).** We investigated whether administration of a TRAIL or a TRAIL- inducer such as an TIC10/ONC201 could suppress tumor growth in the presence of senescent fibroblasts. TIC10/ONC201 or TRAIL was added to the co-culture system. The luciferase reporter assay showed that treatment with TRAIL or ONC201 reduced bioluminescence of HT29-Luc in co-culture in the presence of senescent IMR90 cells as well as in the presence of proliferating IMR90 cells **(Fig. 4A)**. We examined cell viability of the senescent or proliferating IMR90s in response to ONC201 or TRAIL and found that at the doses used, ONC201 did not reduce cell viability of senescent IMR90s, while TRAIL did **(Fig. 4B).** This result agrees with our discovery of the tumor selectivity of ONC201 [20]. Our results suggest that ONC201 suppresses cancer cell growth in the presence of senescent fibroblasts. Furthermore, colony formation assays showed that ONC201 treatment significantly reduced colony formation of HT29 tumor cells in the presence or absence of senescent IMR90 cells **(Fig. 4C).** TRAIL treatment significantly suppressed colony formation of HT29 tumor cells only in the presence of senescent IMR90 cells at the tested doses **(Fig. 4C and 4E).** By contrast, senescent IMR90 fibroblasts conferred cancer cell resistance to 5-Fluorouracil (5-FU), a conventional chemotherapeutic agent used to treat colorectal cancer (CRC). 5-FU treatment significantly reduced colony formation by HT29 CRC cells in the presence of proliferating IMR90 fibroblast cells, with a diminished reduction in the presence of the senescent IMR90 cells **(Fig. 4D-E).** These results agree with previous reports that cancer cells appear to be resistant to 5-FU in the presence of senescent fibroblast cells [21–23].

**Figure 4.**
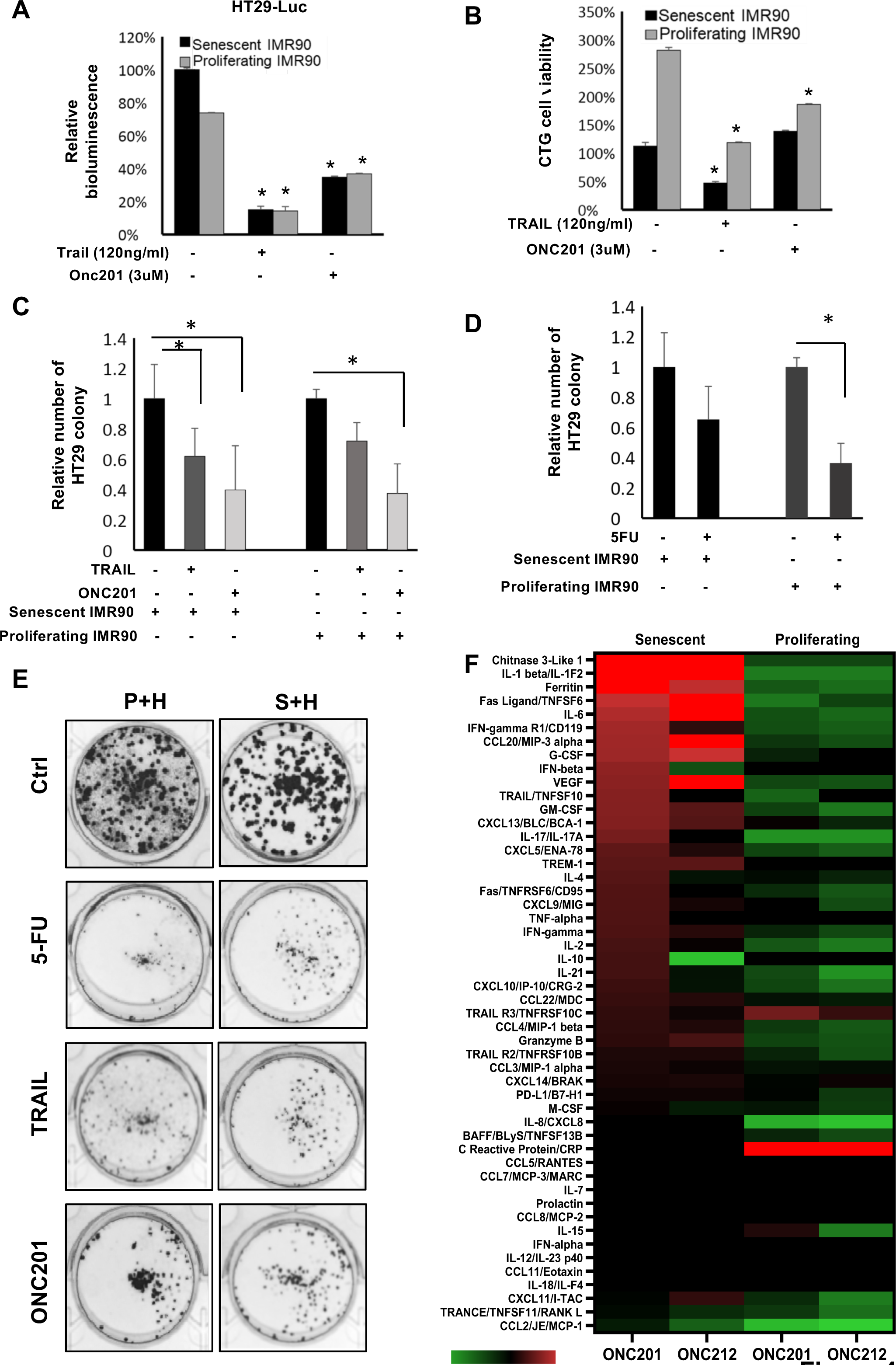
Cancer cell growth modulation in co-culture system with IMR90 fibroblasts in response to different drugs. A. Bioluminescence of HT29-luc cells in two-cell co-culture system treated with TRAIL and TIC10/ONC201 as indicated. Relative bioluminescence was normalized to control for each cohort, respectively. B. Cell viability of IMR90 fibroblasts treated with TRAIL and ONC201. C. HT29 CRC colony formation using co-culture system treated with 3 µM of ONC201 or 100 ng/ml of TRAIL. D. HT29 colony formation in co-culture system treated with 5-FU (1 µg/ml). E. Imaging of HT29 CRC colony formation (quantified in panels C and D). F. Heat map representing cytokine, chemokine, and growth factor profiles. Fold-changes post-treatment with ONC201 (5 µM) or ONC212 (5 µM) are shown for both senescent and proliferating IMR90s. Data are expressed as mean ± SD. *, *p* <0.05. P+H, HT29 cells were cocultured with the proliferating IMR90 cells; S+H, HT29 cells were cocultured with senescent IMR90 cells.

To determine the impact of the Imipridones ONC201 or its potent analogue ONC212 on the cytokinome of both proliferating and senescent cells, we analyzed cell culture supernatants from fibroblasts treated with ONC201 or ONC212. We observed striking differences in response to treatment between senescent and proliferating IMR90 fibroblasts. We observed similar changes with some minor differences in cytokine profiles after treatment with ONC212 as compared with ONC201. Overall, we observed decreased expression of the vast majority of the analytes in response to imipridone treatment of proliferating IMR90s, while we saw increased expression of most analytes in response to treatment of senescent fibroblasts including TRAIL and soluble DR5 **(Fig. 4F).**

### The combination of Imipridones and ABT263 suppresses tumor growth in vivo within a senescent fibroblast reconstructed TME

We explored the impact of clearance of senescent cells in the TRAIL treated TME by combinatorial treatment with ABT263 and ONC201. Colony formation showed that combinatorial treatment with ABT263 and ONC201 reduced colony formation better than either agent alone in different colorectal cancer cells co-cultured with senescent IMR90 (**Fig. 5A and 5B**).

**Figure 5.**
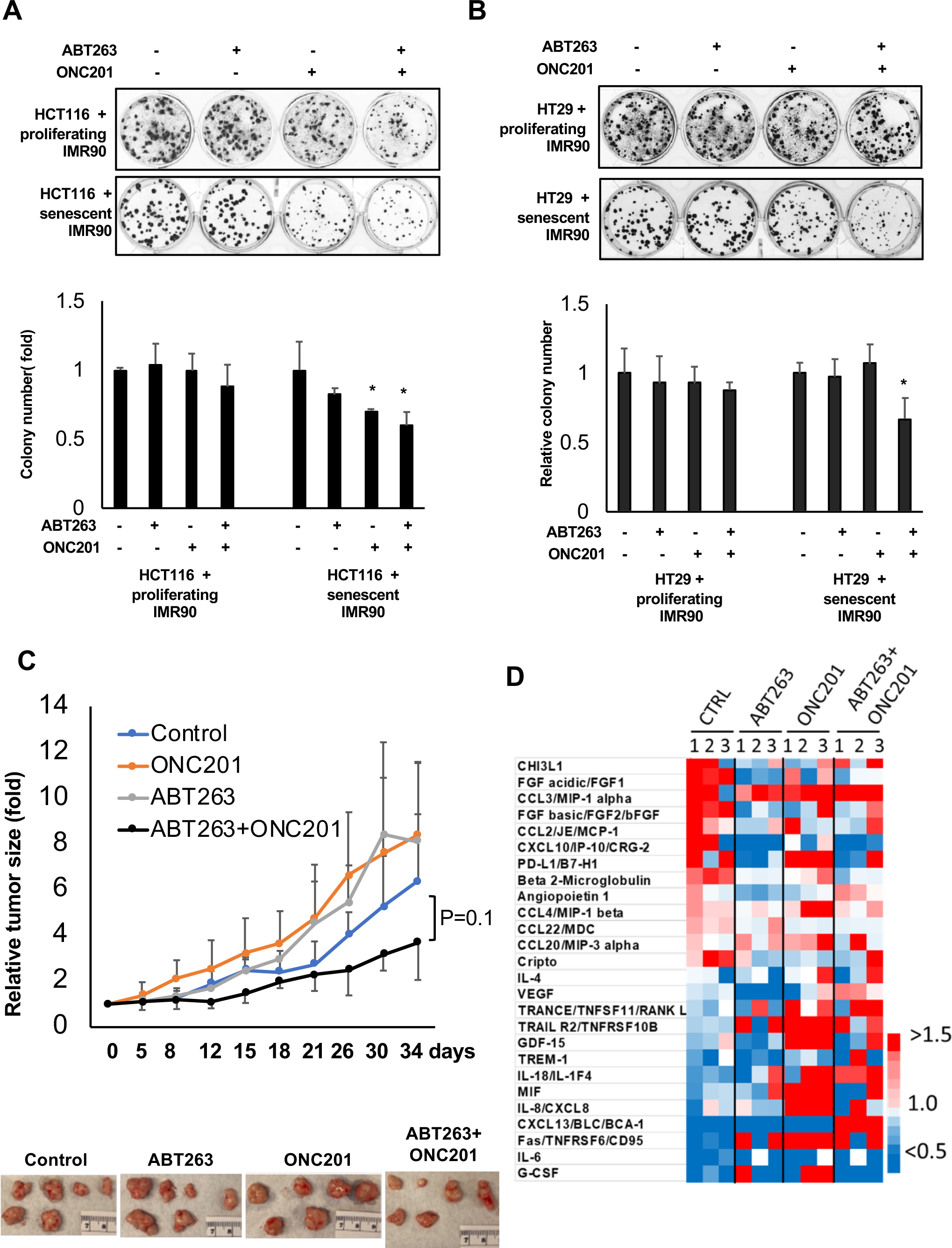
Combinatorial treatment of ABT263 and ONC201 in a senescent reconstructed TME. A. Colony formation assay of HCT116 CRC cells in co-culture system. B. Colony formation assay of HT29 CRC cells in co-culture system. The cells (A and B) were treated with 1 µM of ABT263 and 2 µM of ONC201, or the agents alone. C. Tumor volumes of HT29 tumor xenografts in mice treated with ABT263 and ONC201.D. Heat-map representing fold-changes of serum cytokines in mice with human tumor xenografts containing senescent fibroblasts. Serum samples collected at the end of experiments were subjected to analysis using a human cytokine-profiling panel for bulk cytokine analysis. Relative bioluminescence was normalized to control in each cohort, respectively. Data are expressed as mean ± SD. *, *p* <0.05.

We reconstructed a senescent TME in mice by implanting HT29 CRC cells with senescent fibroblasts in nude mice. The combination of senolytic ABT263 and ONC201 suppressed tumor growth *in vivo* (**Fig. 5C**). Cytokine analysis from mouse blood samples showed that pro-tumorigenic serum cytokines after ABT263 treatment were reduced in mice growing human tumors and senescent fibroblasts in their reconstructed TME (**Fig. 5D**). ONC201 administration increased the TNF superfamily factors including TNFSF11 as compared to senescent cells (**Fig. 5D**) suggesting that ONC201 combined with clearance of senescent IMR90 cells increases innate immune upregulation of TNFSF11 and DR5 secretion. The cytokines induced by individual treatments also appeared in the combination treatment and were correlated with suppression of tumor growth. Collectively, combination of ABT263 and ONC201 depleted senescent CAFs (by ABT263) and by adding extra TRAIL (ONC201 is a TRAIL gene inducer) in the TME. These effects resulted in increased anti-tumor efficacy in the senescent TME.

### Senescent fibroblasts reduce immune-cell mediated tumor cell-cytotoxicity in a tri-culture while imipridones stimulate immune cell death

To investigate the impact of senescent cells on anti-tumor immunity, a tri-culture of natural killer (NK-92) or T cells (TALL-104), senescent or proliferating IMR90 fibroblasts, and HT29 tumor cells was utilized. In the tri-culture with senescent IMR90, HT-29, and NK-92 cells, we observed that NK-92 cells preferentially killed the fibroblasts over the tumor cells at the 8-hour timepoint **(Fig. 6A).** We also observed this preference for natural killer cell-mediated fibroblast killing over tumor killing in the tri-culture containing the proliferating IMR90s **(Fig. 6B).** In our control co-cultures with fibroblasts and tumor cells in the absence of immune cells, we observed minimal cell death due to drug treatment alone with both senescent IMR90s **(Supp.** Fig. 3A) and proliferating IMR90s **(Supp.** Fig. 3B). In the experimental groups containing all three cell types in combination with single drug treatment, we noted increased amounts of natural killer cell-mediated target cell killing with single drug treatments with ONC201 in both senescent IMR90s **(Fig. 6A, 6C)** and proliferating IMR90s **(Fig. 6B, 6D).** ONC201 increased immune cell-mediated target-cell killing as compared to baseline NK-92 cell killing alone **(Fig. 6C-D).** We similarly observed that the immune cells preferentially killed fibroblasts before tumor cells, even in the presence of drug treatment.

**Figure 6.**
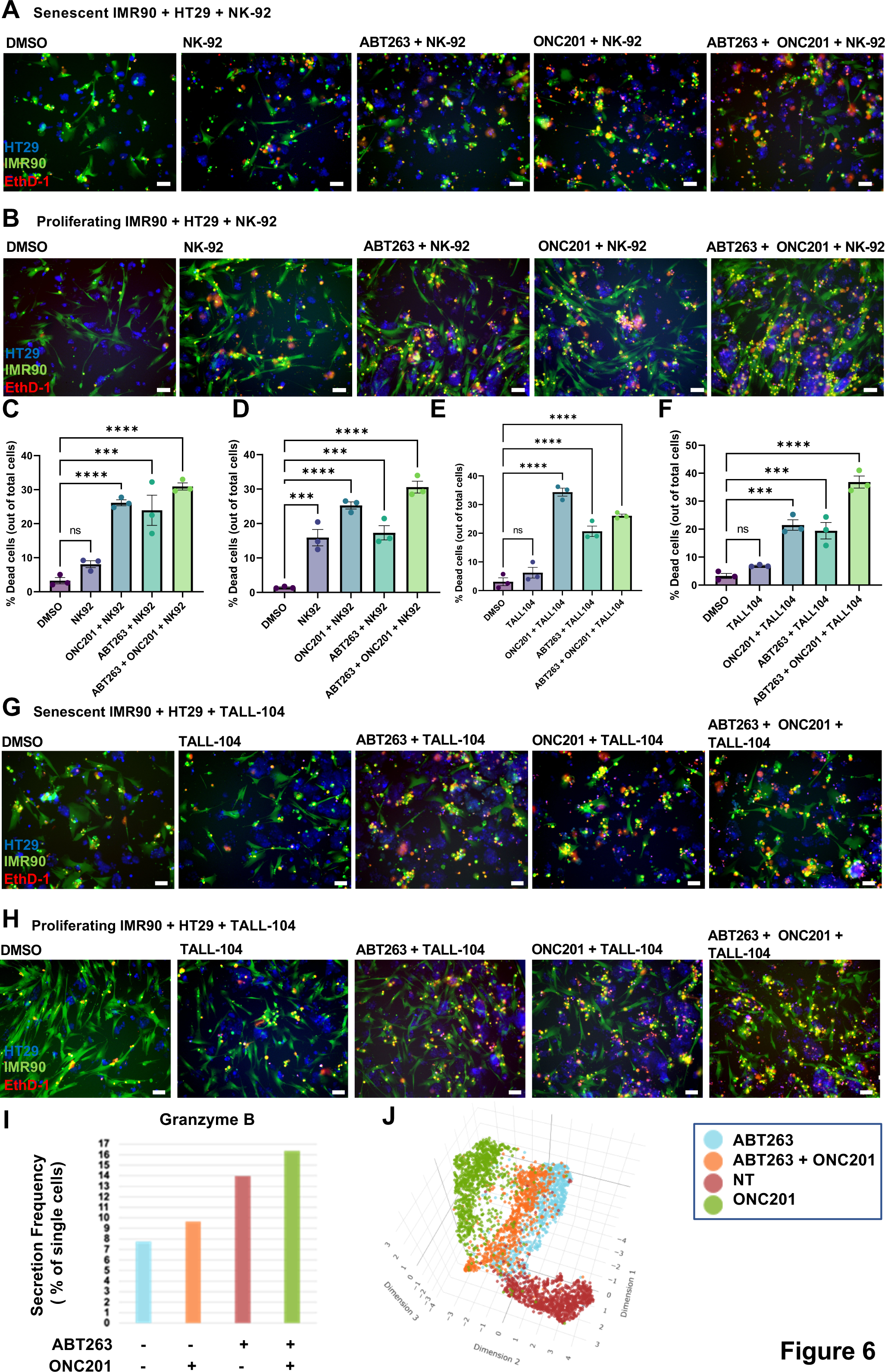
ONC201 increases immune-mediated tumor cell death in a tri-culture assay using both senescent and proliferating fibroblasts. A. Tri-culture of HT29 CRC cells (blue), *senescent* IMR90 fibroblasts (green), and NK-92 cells (unlabeled). B. Tri-culture of HT29 CRC cells (blue), *proliferating* IMR90 fibroblasts (green), and NK-92 cells (unlabeled) at 8-hour post-treatment timepoint. C. Quantification of *senescent* IMR90 + HT29 CRC cells + NK-92 tri-culture (panel A). D. Quantification of *proliferating* IMR90 + HT29 + NK-92 tri-culture (panel B). E. Quantification of *senescent* IMR90 + HT29 + TALL-104 tri-culture (panel G). F. Quantification of *proliferating* IMR90 + HT29 + TALL-104 tri-culture (panel H). G. Tri-culture of HT29 tumor cells (blue), *senescent* IMR90 fibroblasts (green), and TALL-104 T cells (unlabeled). H. Tri-culture of HT29 tumor cells (blue), *proliferating* IMR90 fibroblasts (green), and TALL-104 T cells (unlabeled). Ethidium homodimer (EthD-1) was used to visualize dead cells, 10x magnification, scale bar indicates 100 μm. One-way Anova followed by post-hoc Tukey’s multiple comparisons test was used to determine statistical significance at *p* < 0.05. The following symbols * and ** represent, *p* < 0.05 and *p* < 0.01, respectively. I. TALL-104 cell Granzyme B secretion post-ABT263 or ONC201 treatment was quantified using an IsoPlexis innate-immune chip single-cell cytokine profiling assay. J. A Uniform Manifold Approximation and Projection (UMAP) plot was used to cluster the single TALL-104 cells post-treatment as indicated using IsoPlexis technology. The cells were treated with ABT263 (2 µM) and ONC201 (2 µM), or the agents alone.

Given the preferential natural killer cell-mediated killing we observed, we sought to examine the effect of T-cells in the tri-culture. We added TALL-104 cells to a tri-culture containing senescent or proliferating IMR90s and HT29 CRC cells. ONC201 resulted in the most significant level of target cell death **(Fig. 6E-F).** We noted that T-cells killed fibroblast cells before tumor cells with both senescent and proliferating IMR90s **(Fig. 6G-H)**. Drug treatment of tumor cells and fibroblasts alone resulted in minimal cell death with both senescent and proliferating IMR90s. We noted that treatment of the tri-culture with TIC10/ONC201 stimulated T-cell-mediated tumor cell death **(Fig. 6E-H)**. Single cell cytokine analysis showed that TALL-104 cells had increased secretion of Granzyme B post-treatment with ABT263 and ONC201 **(Fig. 6I).** The secretion frequency of granzyme B (% of single cells) was highest in the combination therapy group including both ABT263 and ONC201. As expected, single TALL-104 cells clustered differently depending on treatment condition when visualized using Uniform Manifold Approximation and Projection (UMAP) clustering **(Fig. 6J).**

## Discussion

Senescence in the TME affects a diversity of components including cancer cells, tumor-associated fibroblasts, and normal epithelial cells. Cellular senescence plays a critical role in tumor formation, progression, relapse, and resistance to therapy. In addition to tumor cell and tumor-associated fibroblast senescence, normal fibroblast senescence appears important to tumor growth and therapy resistance. We demonstrated changes in secreted proteins during the induction of IMR90 fibroblast cell senescence and tested therapeutic strategies such as TRAIL pathway-induction of anti-tumor immunity. The characterization of cytokines profiles in senescent fibroblasts will be useful for guiding cancer therapy strategies as well as future studies involving the senescent TME.

The IMR90 fibroblast represents the normal human cell line most frequently used for senescence research. Our cellular senescence model clearly shows dynamic changes in cytokine profiles with increases in pro-tumor and decreases in anti-tumor cytokines during the induction of senescence in fibroblasts by etoposide. Cytokines that decreased during normal fibroblast senescence are primarily anti-tumorigenic and immunostimulatory. For example, TNF-alpha is known to be a chemoattractant to natural killer and T-cells, activates T cells, and has anti-tumor effects in some contexts [24, 25]. IFN-gamma is an important cytokine that attracts both natural killer- and T-cells and has known anti-tumor effects [26]. IL-12 is a T-cell chemoattractant, and an NK- and T-cell activator [27, 28]. IGFBP-3 upregulates PI3K/Akt/mTOR signaling and downregulates autophagy during cellular aging [29]. Moreover, CCL7/MCP-3/MARC is both a NK- and T-cell chemoattractant and may have anti-tumor effects [30]. Finally, TRAIL/TNFSF10 has anti-tumor effects [31]. The cumulative downregulation of anti-tumorigenic and immuno- stimulatory analytes likely contributes to promotion of cancer in the context of a senescent TME. By contrast, many of the cytokines that increased during the induction of IMR90 cellular senescence were pro-tumorigenic and immunosuppressive. For example, IL-6 is a key factor of the senescent secretome with the ability to promote tumorigenesis and cell proliferation, but can also exert tumor suppressive functions, depending on the cellular context [32]. CXCL10/IP-10 have pro- or anti- tumorigenic effects depending on the context of the cellular signaling [33]. Meanwhile, CCL2/MCP-1 is a known regulatory T-cell chemoattractant with pro-tumorigenic effects [34]. IL-1 alpha induces the production of SASP factors by senescent cells as a result of mTOR activity [35]. G-CSF dampens NK- and T-cell activation and activates regulatory T-cells and has known pro-tumorigenic effects [36]. IL-8/CXCL8, a pro-inflammatory cytokine, is consistently present in the senescent secretome promoting tumor growth [37]. CXCL2/GRO beta/MIP-2/CINC-3, MMP-1, MMP-2, and CXCL1/GRO alpha/KC/CINC-1 are known SASP factors [38]. Finally, IL-1 beta can have pro- or anti-tumor effects depending on the cellular signaling context [39]. The cumulative upregulation of these pro-tumorigenic and immunosuppressive analytes likely contributes to promotion of cancer in the context of a senescent TME.

We demonstrate that changes of cytokine profiles in etoposide-induced fibroblast senescence are regulated by p21(WAF1; CDKN1A). p21 expression also known as senescence-derived inhibitor (sdi1) is one of the hallmarks of senescence induced by DNA damage that results in a permanently arrested cell cycle. A recent study showed that p21 activates the senescence-associated secretory phenotype (SASP) in senescent cells and triggers immune surveillance [40]. We showed that p21-knockdown reverses SASP secretion patterns in senescent IMR90 fibroblasts with an increase in anti-tumor cytokines such as TNFα, ferritin, and IL-18, and a decrease in immunosuppressive inflammatory factors such as CCL2 and IL-8. These data suggest that senescent fibroblasts promote bystander cancer cell growth via suppression of immune surveillance and/or tumor cell apoptosis. Indeed, p21-knockdown in senescent IMR90 fibroblasts suppresses bystander cancer cell growth partially and modestly via TNFα which is a cell death-inducing cytokine [41]. Our results suggest that reduction of anti-tumor cytokine secretion in the TME is a significant mechanism by which senescent fibroblasts promote bystander cancer cell growth. We exploited the knowledge of reduction of TRAIL and DR5 secretion from senescent IMR90 to therapeutically target the senescent TME. As TRAIL inhibits tumor growth in the senescent fibroblast reconstructed tumor microenvironment, targeting the TME by increasing anti-tumor cytokines would contribute to tumor growth inhibition.

Proteins secreted from senescence cells, such as fibroblasts, in the TME may play a role in suppressing the anti-tumor immune response. Indeed, we noted that immune cell-mediated tumor cell-killing was dampened by senescent fibroblasts in co-culture models using both NK- and T-cells. Moreover, ONC201 has been found to increase NK-mediated tumor cell-killing [42]. Our studies show that ONC201 treatment of tri-cultures enhanced immune-mediated tumor cell death in a senescent fibroblast-constructed microenvironment. ONC201 treatment of tumor cells increases secretion of immune activating and recruiting factors such as IFN-α2a, IL-12p70, and IFN-γ–induced protein 10 (IP-10, or CXCL10) [43]. Increased immune stimulatory factors in the TME represents a molecular mechanism for increased immune cell activity post-ONC201 treatment. We noted increased granzyme B secretion frequency in T-cells post-ONC201 treatment. Interestingly, we noted an increase in granzyme B secretion frequency in T-cells post-treatment with senolytic ABT263 and we observed modest increases in immune cell-mediated tumor cell death in tri-cultures treated with ABT263. The differential therapy-induced changes in cytokine profiles by senescent or proliferating cells could impact how tumor cells respond to therapy during cancer treatment.

In summary, we characterized cytokine landscapes to explore modulation of SASP within senescent fibroblasts and sought to promote anti-tumor immunity by inducing TRAIL or by reversing a p21-driven SASP in the senescent TME. Our studies provide strategies for targeting the senescent TME via suppression of bystander cancer cell growth and increased immune cell activation. Moreover, therapeutically-induced changes in the senescent cytokine landscape could help address pro-tumorigenic properties caused by prolonged cellular senescence such during tumor progression and resistance to anti-cancer therapies.

## Acknowledgements

W.S.E-D. is an American Cancer Society Research Professor and is supported by the Mencoff Family University Professorship at Brown University. This work was supported by the Teymour Alireza P’98, P’00 Family Cancer Research Fund established by the Alireza Family. This work was presented in part at the 113th and 114th annual meetings of the American Association for Cancer Research in New Orleans, LA in 2022 and Orlando, FL in 2023, respectively. Figures were created with BioRender.com

## Supplementary Figure Legends

**Supplementary Figure 1.**
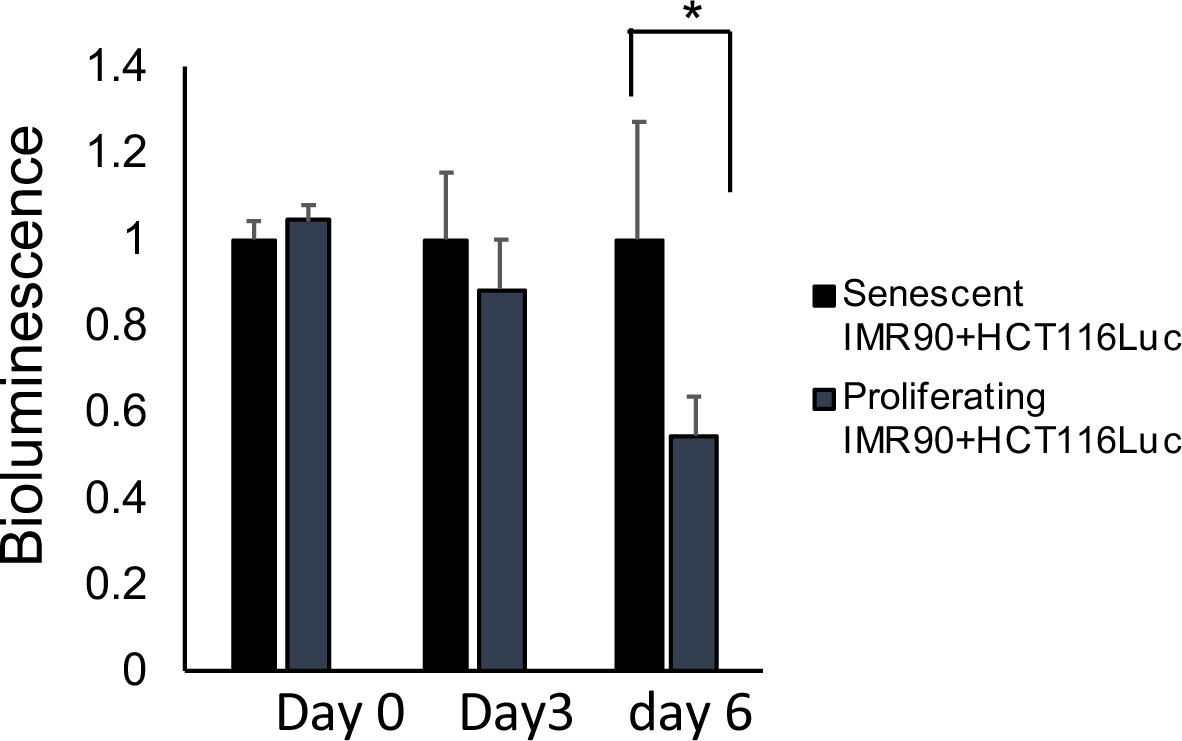
Bioluminescence of cancer cells in the two-cell co-culture system. HCT116-Luc cells were cocultured with the senescent IMR90 or proliferating MR90 cells. Relative bioluminescence was normalized to the control in each cohort, respectively. Data are expressed as mean ± SD. *, *p* <0.05.

**Supplementary Figure 2.**
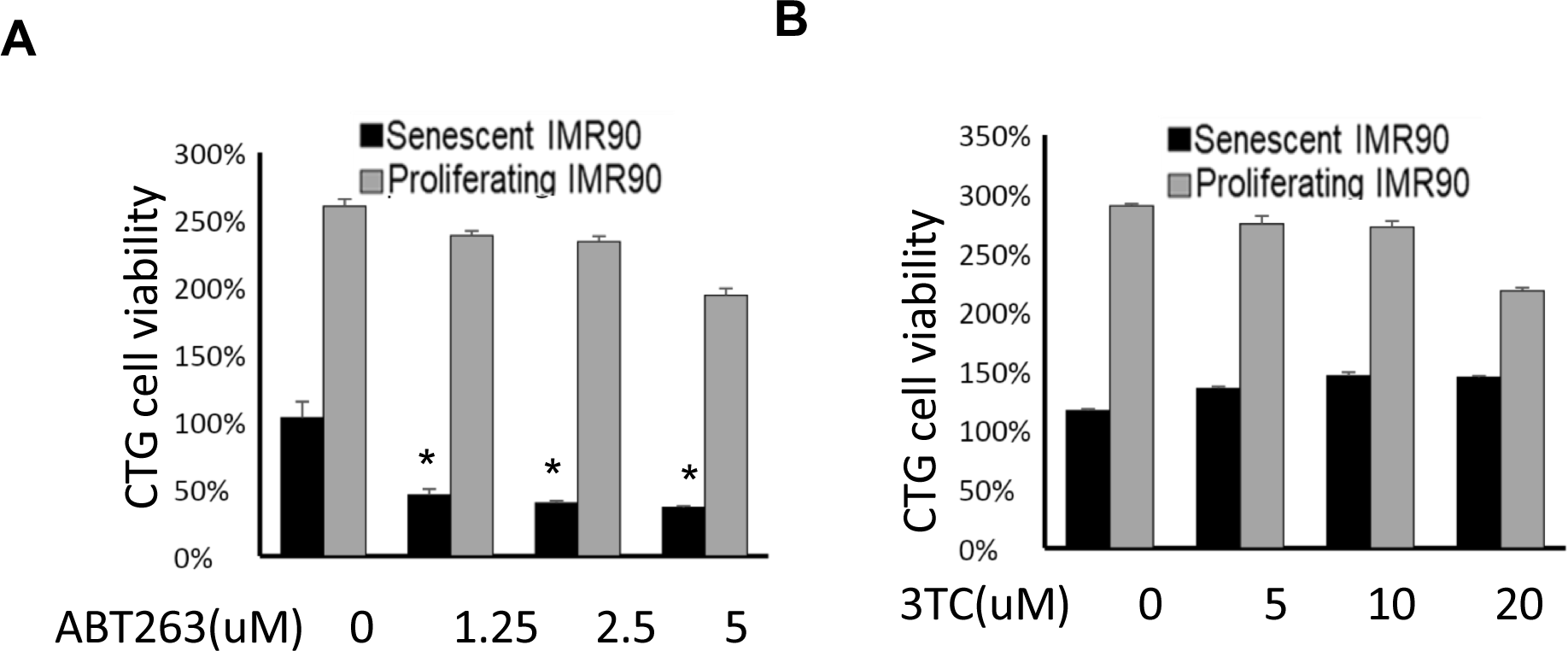
Cell viability. A. Cell viability graph representing senescent and proliferating IMR90 cell viability post treatment with ABT263 at indicated concentrations. B. Cell viability graph representing senescent and proliferating IMR90 cell viability post treatment with 3TC at indicated concentrations.

**Supplementary Figure 3.**
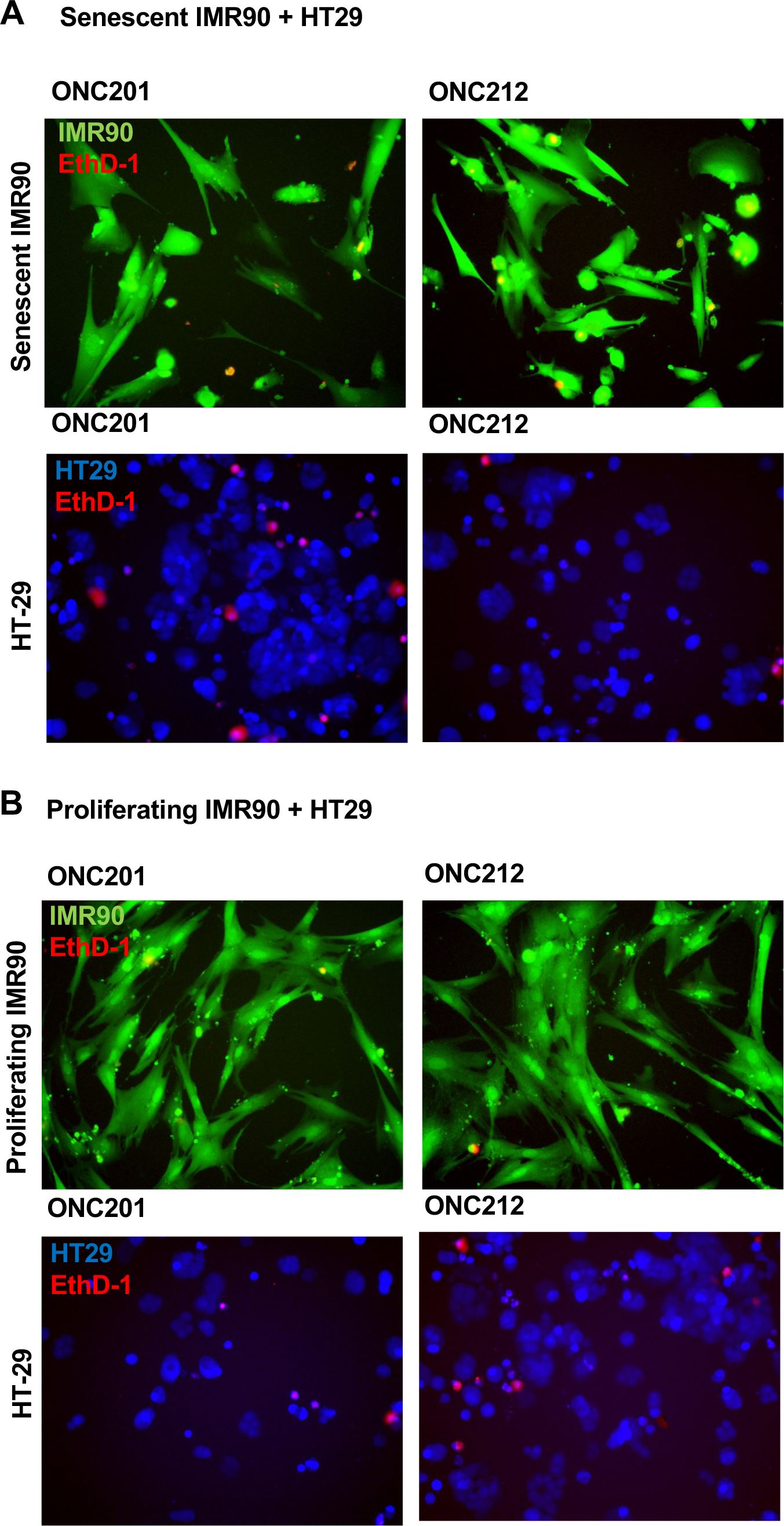
Imagining analysis of cancer cells and IMR90 cells in the two-cell co-culture system. A. Culture of HT29 colorectal cancer cells (blue) or senescent IMR90 fibroblasts (green) with drug treatment controls. B. Culture of HT29 colorectal cancer cells (blue) or proliferating IMR90 fibroblasts (green) with drug treatment controls. 8-hour post-treatment timepoint. Ethidium homodimer (EthD-1) was used to visualize dead cells, 10X magnification.

